# Low levels of neutralizing antibodies against SARS-CoV-2 KP.3.1.1 and XEC in serum from seniors in May 2024

**DOI:** 10.1101/2024.11.18.623742

**Authors:** Even Fossum, Elisabeth Lea Vikse, Anna Hayman Robertson, Asia-Sophia Wolf, Andreas Rohringer, Lill Trogstad, Siri Mjaaland, Olav Hungnes, Karoline Bragstad

**Affiliations:** Department of Virology, Division of Infection Control, Norwegian Institute of Public Health, Oslo, Norway; Department of Method Development and Analytics, Division of Infection Control, Norwegian Institute of Public Health, Oslo, Norway

## Abstract

New immune evasive variants of SARS-CoV-2 may increase infections and hospitalizations in risk groups, such as the elderly. In this study we evaluated neutralizing antibodies against KP.3.1.1 and XEC, virus variants that are either widely distributed or on the rise globally, in sera from a cohort of seniors aged 68 - 82 years from April/May 2024. Neutralizing responses were low against both KP.3.1.1 and XEC, supporting the recommendation of an updated covid-19 vaccine booster in this age group.

## Background

Vaccination and infections have contributed to a high level of immunity against SARS-CoV-2 in the Norwegian population (1). Nevertheless, new immune evasive SARS-CoV-2 variants continue to emerge and cause periodic increases in infections, hospitalizations and deaths, especially in risk groups such as the elderly (2). August 2023 saw the emergence of the BA.2.86 variant which differed from the then circulating XBB derived EG.5.1 strain by >30 mutations in the Spike protein (3, 4). In Norway, BA.2.86 derived JN.1-variants became dominant in October/November 2023 (Figure 1A). Infections with the JN.1-derived KP.2 variants, which were defined by addition of the so called “FLiRT”-mutations (F456L and R346T), started to increase from March 2024(5). KP.2-variants were from June 2024 overtaken by KP.3-variants containing the “FLuQE”-mutations (F456L and Q493E), and since July the KP.3.1.1 variant containing the S31 deletion have dominated (6). Recently, the prevalence of the XEC-variant (recombination of BA.2.86 derived variants KS.1 and KP.3.3) has started to rise, with recent studies suggesting greater immune evasion compared to KP.3.1.1 (7, 8). To evaluate existing humoral immunity in the elderly prior to the winter season 2024/25, we performed neutralization assays against KP.3.1.1 and XEC on serum samples harvested from seniors in April/ May 2024.

**Figure 1:**
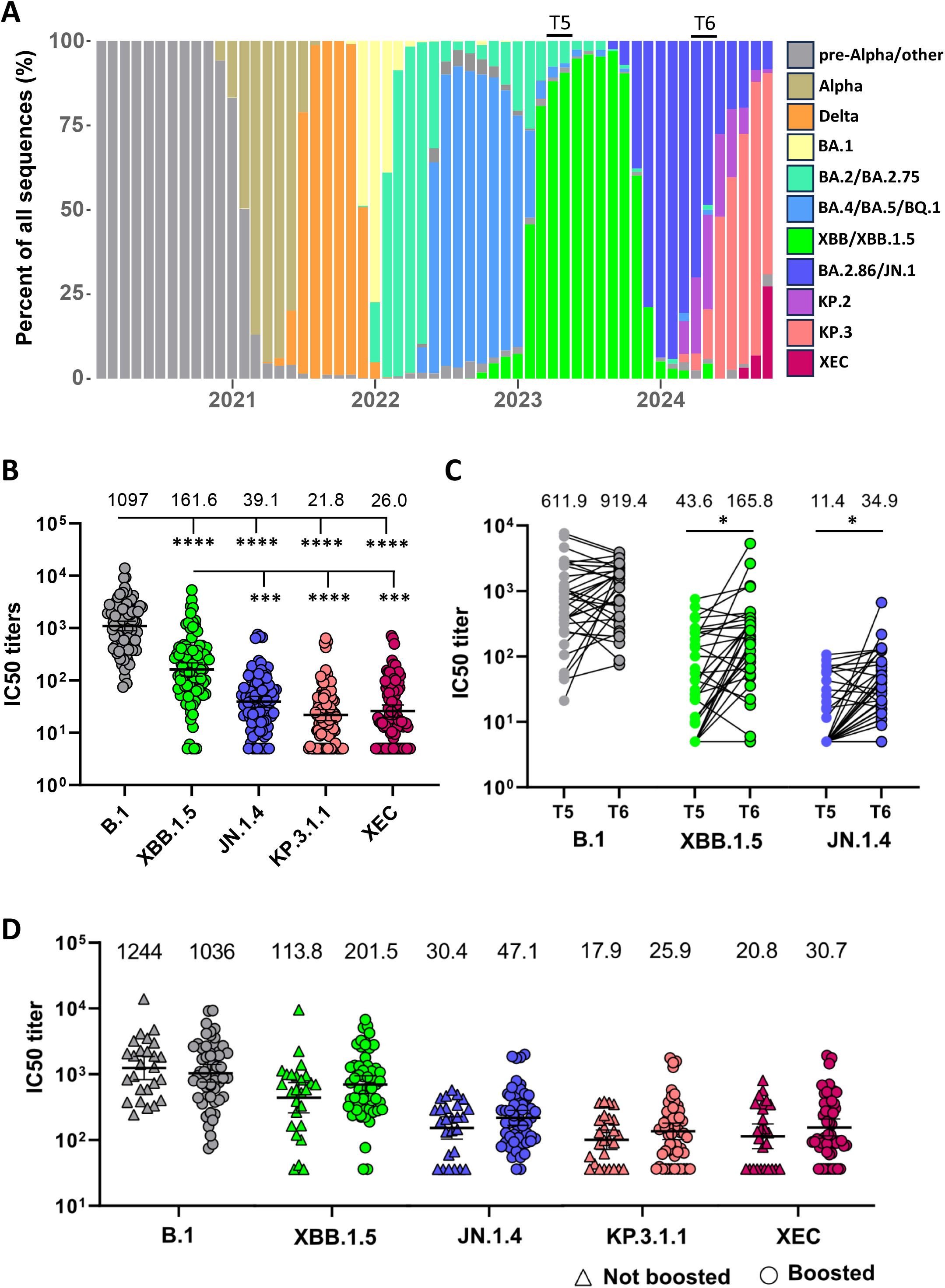
Neutralizing titers against SARS-CoV-2 variants in sera from seniors. **A)** Prevalence of SARS-CoV-2 variants in Norway determined by whole-genome sequencing from January 2020 to October 2024. T5 and T6 indicate time points for serum sampling in April/May 2023 and April/May 2024, respectively. **B)** Neutralizing titers in seniors (n = 97) against B.1, XBB.1.5, JN.1.4, KP.3.1.1 and XEC in serum samples collected in April/May 2024. The number above each column indicates geometric mean titer. **C)** Change in neutralizing titers against B.1, XBB.1.5 and JN.1.4 in 33 individuals who donated sera at time points T5 and T6. **D)** Serum samples from April/May 2024 were divided into two groups depending on whether the donor reported to have received the monovalent XBB.1.5 vaccine, excluding individuals with reported infection between time points T5 and T6. In total, 55 donors had received an XBB.1.5 booster dose and 26 donors had not. The number above each column indicates geometric mean titer. The data presented in panels **B** – **D** are IC50 neutralizing titers, with geometric mean ± 95% CI shown in panel **B** and **D**. Significant difference analyzed in **B)** by one-way ANOVA with Tukey`s multiple comparison test, in **C)** by Paired t-test and in **D)** by Mann-Whitney test. * = p < 0.05, *** = p < 0.001 and **** = p < 0.0001.

### Study cohort and experimental analysis

The Senior Cohort was established towards the end of 2020 to study the consequences of COVID-19 disease and the effects and safety of COVID-19 vaccines in elderly individuals. Approx. 5000 individuals aged 65 – 80 years living in Oslo, Norway, consented to participate in answering regular questionnaires, and approx. 500 individuals consented to donating longitudinal blood samples, as previously described (9). Of these, 97 individuals donated blood in April/May 2024. Information on individual vaccination and infection status was obtained through regular questionnaires from May 2021 up to the time of serum donation, in addition through the Norwegian Immunization Registry (SYSVAK) and the Norwegian Surveillance System for Communicable Disease (MSIS).

SARS-CoV-2 variants were isolated from clinical samples submitted for surveillance to the Norwegian Institute of Public Health (NIPH) in VeroE6-TMPRSS2 cells as previously described (10). JN.1.4 (available in GISAID EpiCoV with accession number EPI_ISL_18735198), KP.3.1.1 (EPI_ISL_19268292) and XEC (EPI_ISL_19393351) were passaged twice in VeroE6 before titration and assessed in a live virus neutralization assay as previously described (10). SARS-CoV-2 variants B.1 (EPI_ISL_449791) and XBB.1.5 (EPI_ISL_16969674) were previously isolated (1).

### Reduced neutralizing titers against KP.3.1.1 and XEC

To evaluate immunity in seniors in Norway against recent SARS-CoV-2 variants, 97 serum samples were harvested from individuals aged 68 - 82 years in April/May 2024. At the time, BA.2.86 derived JN.1 variants were dominant in Norway, although infections with BA.2.86 derived KP.2 and KP.3 variants were increasing (Figure 1A). All individuals had received at least three doses of mRNA vaccine (range 3 – 6 doses with a median of 5 doses), and 59 (60.8%) individuals had reported at least one previous breakthrough infections, thus reflecting the heterogeneity of immunity currently present in the Norwegian population. The serum samples were analyzed in a live-virus neutralization assay against SARS-CoV-2 variants B.1, XBB.1.5, JN.1.4, KP.3.1.1 and XEC (Figure 1B).

While neutralizing titers against B.1 were high, there was a significant reduction in neutralization of XBB.1.5, JN.1.4, KP.3.1.1 and XEC. Neutralizing titers against XBB.1.5 were also significantly higher than JN.1.4, KP.3.1.1 and XEC. (Figure 1B). Neutralizing titers against JN.1.4 were higher than KP.3.1.1 and XEC, but the difference was not significant (Figure 1B). For KP.3.1.1 and XEC the levels of neutralizing titers were very low. We did not see a further reduction for XEC compared to KP.3.1.1 as has been reported in pseudo-neutralization assays (7, 8).

Of the 97 individuals that donated serum in April/May 2024 (“T6”), 33 individuals had also donated sera a year earlier in April/May 2023 (“T5”). Interestingly, the level of neutralization for B.1 remained constant from one year to the next (Figure 1C), which could reflect continuous back-boosting of B.1 neutralizing responses through infections and/or vaccination. For both XBB.1.5 and JN.1.4 there was a significant increase in neutralizing antibodies from 2023 to 2024 potentially reflecting administration of the monovalent XBB.1.5 vaccine in the fall 2023 and circulation of the XBB.1.5 derived variants throughout 2023 and JN.1 derived variants from November 2023 (Figure 1A and C).

To better assess the effects of the XBB.1.5 booster vaccine on neutralizing antibody responses, serum samples collected in April/May 2024 were divided into boosted and un-boosted based on vaccination in October/November 2023. Individuals that had reported a positive SARS-CoV-2 tested between time points T5 and T6 were excluded from the analysis. While boosted individuals did have higher neutralizing titers against XBB.1.5, JN.1.4, KP.3.1.1 and XEC, the difference was not significant for any of the variants (Figure 1D).

## Discussion

While neutralizing antibodies (NAb) are recognized as a correlate of protection against SARS-CoV-2 (11, 12), it is less clear what constitutes a protective level of NAb. The significant reduction in antibody titer from B.1 and XBB.1.5 to KP.3.1.1 and XEC does, however, indicate that the seniors have limited protection against current strains. Our results also indicate that prior immunization with the XBB.1.5 monovalent booster in the fall of 2023 is unlikely to protect against infection with the KP.3.1.1 and XEC strains in the fall/winter of 2024, which is in accordance with a recent study showing limited protection from infection >12 weeks after administration of the XBB.1.5 vaccine (13). However, immunization with an updated JN.1 or KP.2 monovalent vaccine has been reported to increase neutralizing antibodies against the KP.3.1.1 and XEC variants (14). A concern is that only 54% of individuals in Norway aged 65 years or older took a SARS-CoV-2 vaccine during the 2023/24 season (15), highlighting a need to better inform this age group of the benefits of boosting immune responses with an updated SARS-CoV-2 vaccine.

A limitation of our study is that data on vaccinations from July 2023 and on infection from January 2022 were obtained through self-reporting, and we can therefore not rule out errors or underreporting. However, previous data from the same individuals on self-reported immunization and infection have been highly consistent with data from the health registries (SYSVAK and MSIS). It should also be noted that our study only evaluates neutralizing antibodies against recent strains and does not consider the role of T cells. Indeed, previous studies with the same cohort have indicated that T cell responses are more stable against new virus variants (9).

## Conclusion

Our study indicates that seniors have very low levels of neutralizing antibodies against recent SARS-CoV-2 strains KP.3.1.1 and XEC. Antibodies induced by previous vaccination with an XBB.1.5 booster in the fall of 2023 are unlikely to protect against new infections in the winter of 2024/25. The results support current recommendation of an updated booster covid-19 vaccines for this age-group.

## Ethics statement

Blood samples and medical information were collected with written consent from all participants. The study was approved by the Regional Committees for Medical and Health Research Ethics Southeast (REK) with reference nr. 229359. All data were pseudo anonymized to protect patient privacy and confidentiality.

## Data availability

Anonymized data can be shared in accordance with the data sharing policy of NIPH.

## Acknowledgements

We would like to thank the participants in the Senior Cohort who donated blood samples. We would also like to thank the national microbiological laboratories in Norway for providing clinical samples as part of the national COVID-19 surveillance, allowing us to isolate the latest SARS-CoV-2 variants in order to investigate immunity towards these. We thank Rasmus Kopperud Riis for sequence analysis of virus isolates, and all our highly skilled colleagues at the Department of Virology that have made this study possible.

## Authors contribution

E.F. and E.L.V. performed the experiments and analyzed results, A.H.R., A.S.W, L.T. and S.M. collected biological samples, data from questionnaires and established and coordinated the cohort, A.R. collected analyzed SARS-CoV-2 surveillance data, O.H. and K.B. supervised the study. All authors assisted in the design and conceptualization of the study. E.F. wrote first draft of the manuscript, while all authors revised and approved the final version.

